# Sphingosine 1-phosphate activates the MAP3K1-JNK pathway to promote epithelial movement and morphogenesis

**DOI:** 10.1101/2020.06.23.167304

**Authors:** Jingjing Wang, Maureen Mongan, Jerold Chun, Ying Xia

## Abstract

MAP 3 kinase 1 (MAP3K1) plays an essential role in embryonic eyelid development. It regulates epithelial morphogenesis through the spatial-temporal activation of Jun N-terminal kinases (JNKs), resulting in forward progression of the embryonic eyelid epithelial cells to enable eyelid closure. The developmental signals that activate the MAP3K1-JNK pathway are still unknown, mainly due to the lack of suitable keratinocyte lines to elucidate the mechanisms of pathway regulation. To address this deficiency, we developed a straightforward method for long-term culture of mouse keratinocytes in feeder-free conditions using Ca^2+^-free media. Cells grown under these conditions displayed characteristic basal epithelial morphology and keratin 14 expression, but did not form tight- or adherens-junctions. Increased extracellular Ca^2+^ levels restored the formation of cell-cell junctions. Using keratinocyte lines derived from wild type and *Map3k1*-deficient mice, we found that sphingosine 1-phosphate (S1P) activated the JNK-c-JUN pathways in a manner dependent on MAP3K1 kinase activity and that this MAP3K1-mediated signaling led to epithelial cell migration. The *in vivo* roles of this pathway were examined through crossing of genetic mutant mice. Loss-of-function of the S1P receptor *(S1pr) 2/*3 became haploinsufficient only when combined with *Map3k1* and *Jnk1* mutations such that the compound mutants displayed eyelid closure defects, suggesting these gene products cooperated in eye morphogenesis. Results of this work establish the S1PR-MAP3K1-JNK pathway as a crucial signaling mechanism for epithelial cell movement and morphogenesis.

## Introduction

Signal transduction through the mitogen-activated protein kinase (MAPK) pathways is a central mechanism that regulates cellular responses to extracellular stimuli (1). A typical MAPK pathway is comprised of a MAP3K, a MAP2K and a MAPK, in which the MAP3Ks define tissue-, cell type- and stimuli-specificity in MAPK activity (2). The MAP3K1, a serine/threonine protein kinase and a member of the MAP3K superfamily, plays diverse roles in immune system development and function, injury repair, vasculature remodeling and tumor progression (3-10). Additionally, studies in independent genetic mouse models reveal an essential role of MAP3K1 in eye development (11). Mice lacking the full-length or the kinase domain of MAP3K1, or those harboring a spontaneous mutation where a 27.5-kb deletion on chromosome 13 resulting in the elimination of eight exons of the *Map3k1* gene, exhibit an eye-open at birth (EOB) phenotype (12-14). These genetic data indicate that MAP3K1 and its kinase activity are essential for eye development.

The EOB defect is the result of impaired embryonic eyelid closure, a morphogenetic event common in mammalian eye development (15). In mouse embryogenesis, the opposing eyelids move forward and ultimately meet and fuse at embryonic day (E) 15 - E16, resulting in a closed eyelid covering the ocular surface (16-18). The eyelids remain closed at birth, serving as a protective barrier for the immature eye; they re-open at postnatal day (P) 14. The failure of eyelid closure is not life-threatening and the resultant phenotype is easy to detect, enabling the identification of the EOB defects in at least 160 genetic mutant strains (http://www.informatics.jax.org/mp/annotations/MP:0001302). These mutant strains underscore the genetic complexity behind eyelid morphogenesis (11,19). Since embryonic eyelid closure is driven by epithelial forward movement, the EOB mice have become convenient tools to delineate the molecular and signaling mechanisms of epithelial morphogenesis (20-23).

In epithelial cells of the embryonic eyelids, MAP3K1 mediates the phosphorylation and activation of the MKK4 MAP2Ks, which in turn activate the Jun N-terminimal kinase (JNK) MAPKs (12,13,24-26). Molecular studies in cultured cells and genetic mutant mice have further identified RhoA, a small GTPase, as one of the upstream activators of the MAP3K1-JNK pathway (27,28). A potent RhoA activator is sphingosine 1-phosphate (S1P), a phospholipid signaling molecule (29,30). S1P is present abundantly in the circulation and specific tissues; it binds to and activates the S1P receptors (S1PRs), which are G protein coupled receptors (GPCRs) that in turn transmit the signals intracellularly to elicit biological responses (31-35). The S1P/S1PR signal is essential for embryonic eyelid closure, because simultaneous deletion of *S1pr2* and *S1pr3* genes leads to the EOB phenotype similar to that found in the *Map3k1*-null pups (36). The relationships between S1P and MAP3K1 signaling in eyelid development, however, are yet to be investigated.

While keratinocytes derived from genetic mutant mice are valuable tools to investigate the mechanisms of epithelial abnormalities, methods have been limited mostly to the use of primary cultures of murine keratinocytes (37-39). Here we report the long-term growth of murine epithelial cells in low-Ca^2+^ keratinocyte serum-free medium (KSFM) without the requirement of serum and feeder cells. The medium permitted the growth of a few clones to reach confluence. Cells derived from these clones can be subcultured for >60 passages, they express proliferation and basal keratinocyte markers, and can be induced to establish cell-cell junctions. The cells can also be stored in liquid nitrogen and recovered, and retain the same growth and passaging capacity of the parent clones. With this method, we derived cells from wild type and *Map3k1*-deficient mice and obtained an essentially endless supply of genetically-modified keratinocytes. These cells, together with genetic mutant mice, enabled the identification of S1P as the physiological stimulus that activates the MAP3K1-JNK pathway, leading to epithelial movement and embryonic eyelid closure.

## Results

### Long-term mouse keratinocyte culture in media with low extracellular calcium

Three commercial media: (i), the KSFM without Ca^2+^ (KSFM-Ca^2+^), (ii), the KSFM with Ca^2+^ (KSFM+Ca^2+^), and (iii), the defined KSFM (DKSFM), were evaluated in mouse keratinocyte culture. The KSFM+Ca^2+^ and KSFM-Ca^2+^, containing the same growth factor supplements, were different only in Ca^2+^ concentration. The KSFM-Ca^2+^ contained trace amounts of calcium (0.06 mM), a level ∼60% lower than the ∼0.15 mM levels in KSFM+Ca^2+^ and DKSFM (Fig. 1A).

**Figure 1.**
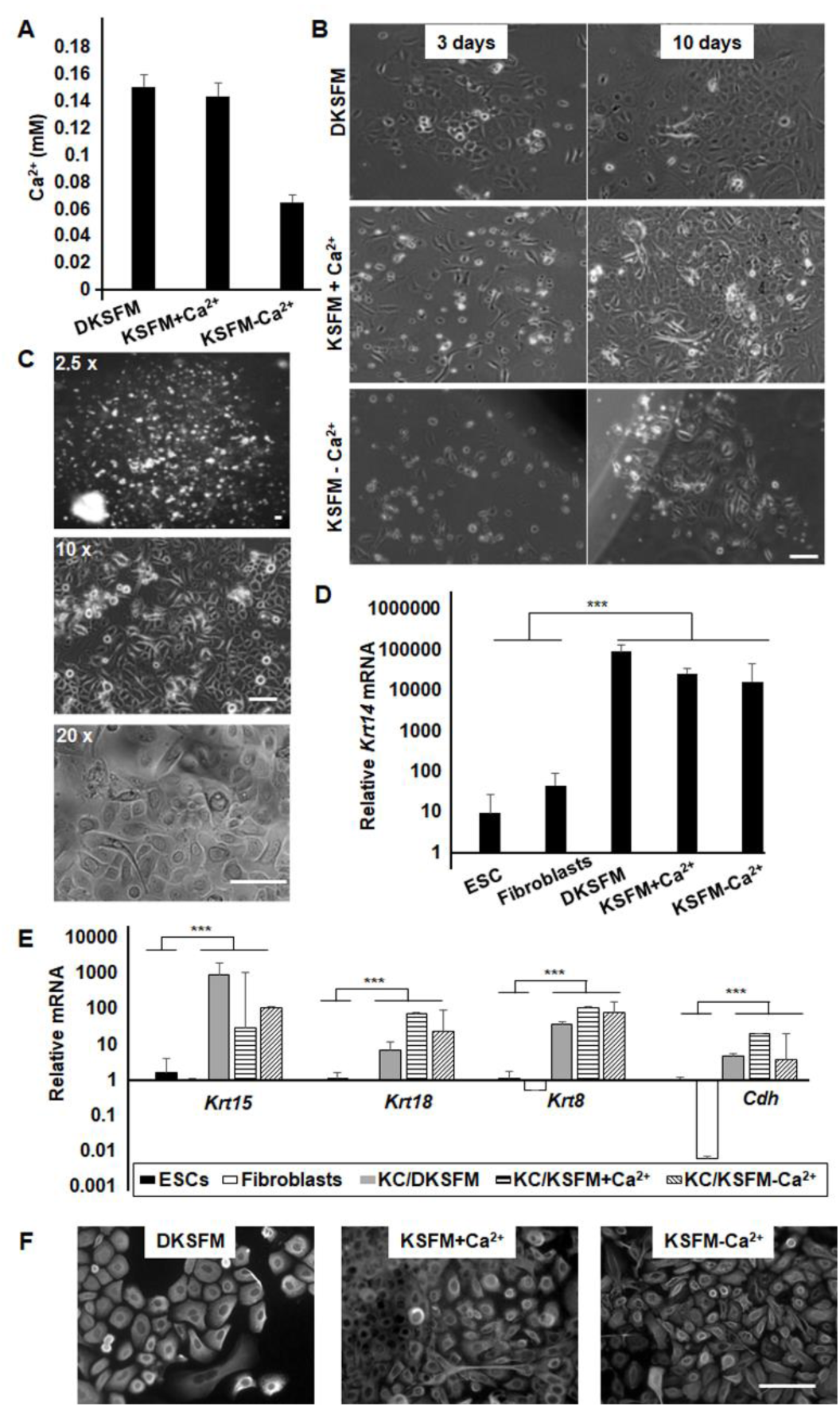
Culture conditions of the mouse keratinocytes. (A) The calcium concentrations in three commercial keratinocyte growth media. (B) Bright-field microscopic images of cells grown for 3 or 10 days in culture. (C) Images of colonies of cells grown in KSFM-Ca^2+^ medium for an extended period of time. The expression levels of keratinocyte-specific genes (D) *Krt14* and (E) *Krt8, Krt15, Krt18* and *Cdh*, were determined by RT-PCR. Expression in cells grown in the three types of KSFM medium was compared to that in ESCs and fibroblasts. (F) Cells were subjected to immunofluorescence staining with anti-K14 and photographed under fluorescent microscope. Results represent data from at least three duplicate experiments +/-SD, and *** p<0.001 was considered significant. The scale bars in microscope image correspond to 100 µm.

The keratinocytes isolated fresh from newborn pups were attached well on collagen IV-coated plates, reproduced quickly and became confluent in 1-2 weeks in DKSFM and KSFM+Ca^2+^ media. A majority of the cells displayed similar morphologies with regular and polygonal shapes (Fig. 1B). The cells grown in KSFM-Ca^2+^ medium, on the other hand, were relatively inefficient in attachment and growth; however, a few small cell clusters became visible by 1-2 weeks in culture (Fig. 1C). Some of these clusters continued to grow, forming large colonies in 1-2 months, and ultimately growing to confluence in a few months. These cells also displayed epithelial morphologies with polygonal shapes and regular dimensions. The colony-forming efficiency varied from batch to batch with an average efficiency of approximately 0.002%.

Cells grown in either KSFM+Ca^2+^ or DKSFM were not easily amplified through passaging, consistent with the recognized challenges in mouse keratinocyte subculture (40-43). Cells in KSFM-Ca^2+^, in contrast, could be extensively amplified through passaging, up to 60 times, resulting in a >10^30^-fold increase in cell number, and the longest culture was maintained up to 18 months. Following storage in liquid nitrogen, the cells retained the growth and expansion capacities.

The identity of these cells was characterized by the examination of keratinocyte-specific gene expression. The expression of *Krt14*, the basal keratinocyte gene, as well as other keratinocyte markers, such as *Krt15, Krt 8, Krt 18* and *Cdh*, was robust in cells grown in KSFM-Ca^2+^, KSFM+Ca^2+^ and DKSFM (Figs. 1D and 1E). In contrast, the expression of these genes was almost undetectable in embryonic stem cells (ESCs) and fibroblasts. Immunostaining showed that cells grown in the different KSFM media exhibited similar K14 expression in cytosol and perinuclei (Fig. 1F). Hence, all three media support epidermal keratinocyte culture, but only the KSFM-Ca^2+^ preferentially supports keratinocyte with long-term proliferative output.

### Molecular characteristics of keratinocytes in long-term culture

The proliferation potential of cells grown in KSFM-Ca^2+^ was assessed by colony-forming efficiency (CFE) assays. The CFE varied between lines derived from mice of the same genetic background, ranging from 0.5 - 6% (N=6) (Table 1). The colonies displayed different shapes and sizes, and there was a notable correlation between CFE and the number of holoclone-like colonies, exhibiting uniform, round shapes, and compact cell contents (44)(Fig. 2A). All 6 lines had similar growth rates with an average doubling time of 5.2 +/- 1.2 days (Fig. 2B). The doubling time decreased as the passage number increased (Fig. 2C and Fig. S1).

**Table 1.** Cultured Murine Epithelial Lines

| **Cell lines** | **Colony forming efficiency (%)** |
| --- | --- |
| 1 | 0.48 |
| 2 | 0.69 |
| 3 | 1.4 |
| 4 | 2.9 |
| 5 | 3.75 |
| 6 | 5.93 |

**Figure 2.**
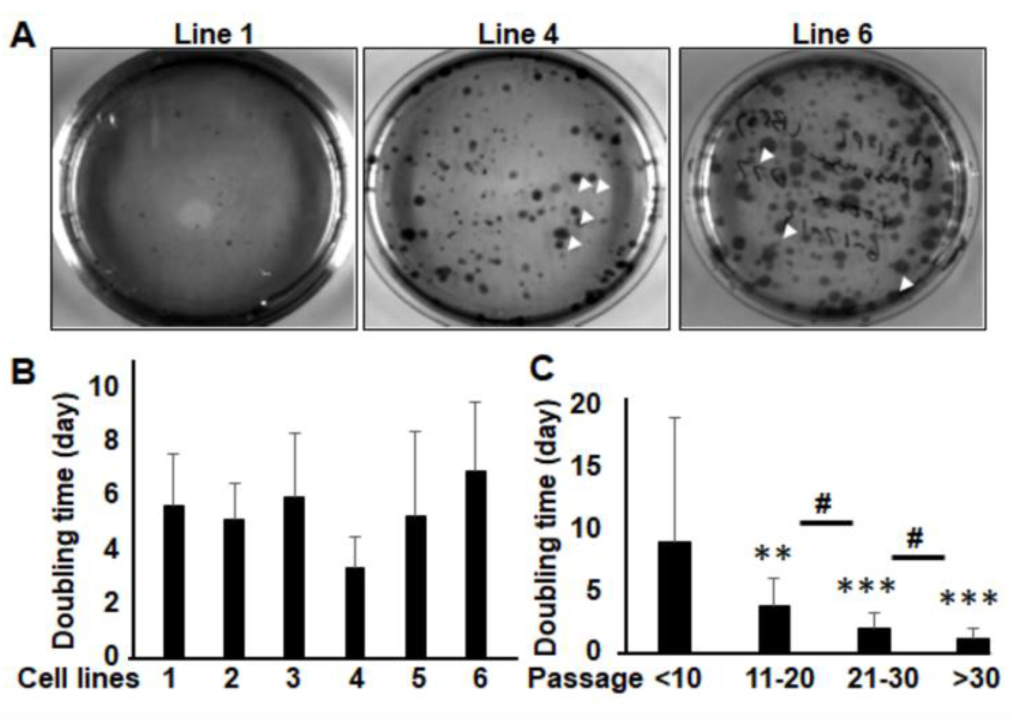
The growth properties of long-term keratinocyte culture. (A) Representative photos of colonies formed with different keratinocyte lines derived in KSFM-Ca^2+^, with varied colony-forming efficiency. Arrowheads indicate at representative holoclones. The average doubling time of (B) each cell line, and (C) all lines at different passages, calculated based on the cell counts upon passaging and days of growth +/-SD. The doubling time of medium and high passages was significantly reduced relative to that of low passages (**p<0.01 and ***p<0.001). Similarly, cells at >30 passages grew much faster than at 21-30 passages, which in turn grew faster than at 11-20 passages (# p<0.01).

The sustained keratinocyte growth could be due to the selective enrichment of epidermal stem cells (45-48). We evaluated this possibility by examining the expression of epidermal stem cell markers and found that the expression of *Cd34, Krt15, Krt19* and *Sca-1* was not much different in cells grown in any of the three media (Fig. 3A). On the other hand, the expression of *Krt10*, a marker of outer-layer terminal epidermal differentiation, and *ColIa1*, encoding extracellular matrix proteins of the epidermis, were markedly less abundant in the KSFM-Ca^2+^ cells than the KSFM+Ca^2+^ and DKSFM cells (Fig. 3B). Conversely, the expression of α6 integrin, a marker of long-term proliferative potential, and *Tp63*, a transcription factor and a marker for proliferative epithelial cells, was more abundant in the KSFM-Ca^2+^ cells (49-51) (Fig. 3C).

**Figure 3.**
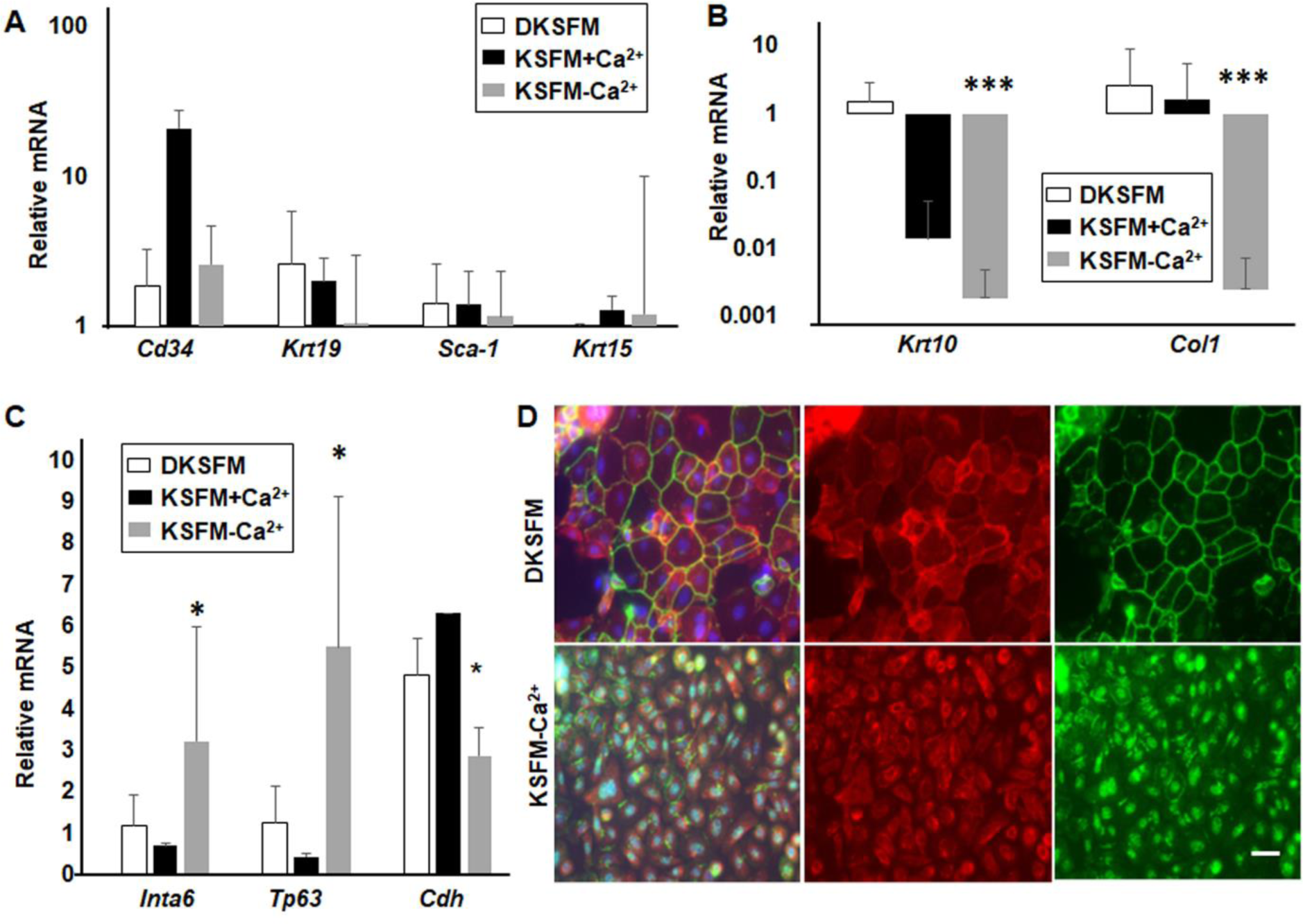
The molecular characteristics of long-term keratinocyte culture. mRNA isolated from primary keratinocytes in DKSFM and KSFM+Ca^2+^, as well as long-term cultured keratinocytes in KSFM-Ca^2+^, was examined by RT-PCR for expression of markers of: (A) epidermal stem cells and progenitors (*Cd34, Krt19, Sca-1* and *Krt15*), (B) epidermal terminal differentiation (*Krt10* and *ColI*), and (C) epithelial cell proliferation (*Inta6* and *Tp63*) and cell-cell junctions (*Cdh*). (D) Immunofluorescence staining for E-Cadherin and ZO-1, which were located on plasma membrane at cell-cell contacts in cells grown in DKSFM, but were mostly detected in the nucleus and/or cytosol in the long-term keratinocyte in KSFM-Ca^2+^. The scale bars in microscope images correspond to 100 µm. Error bars represent Mean +/- SD *p<0.05 and ***p<0.001 are considered significantly different from values in cells grown in DKSFM.

Cells grown in the KSFM-Ca^2+^ also had reduced expression of *Cdh*, which encodes the E-Cadherin that mediates the formation of adherens junctions in basal keratinocytes (52) (Fig. 3C). Correspondingly, adherens junctions were detected by immunostaining in cells grown in DKSFM but not in KSFM-Ca^2+^ (Fig. 3D). The Zonula occludens-1 (ZO-1), component of the tight junctions that provide semipermeable barrier for ions and solutes of the epithelial tissues, was also present at the plasma membrane of cells in DKSFM, but absent in cells in KSFM-Ca^2+^ (Fig. 3D). To determine if higher extracellular calcium level were responsible for the formation of cell-cell junctions, we switched the growth medium from KSFM-Ca^2+^ to either KSFM+Ca^2+^ or DKSFM, resulting in an increase in calcium from 0.06 mM to 0.15 mM. Growing in media with higher calcium for 2 days was sufficient to increase E-Cadherin expression and induce the formation of adherens and tight junctions (Figs. 4A and 4B).

**Figure 4.**
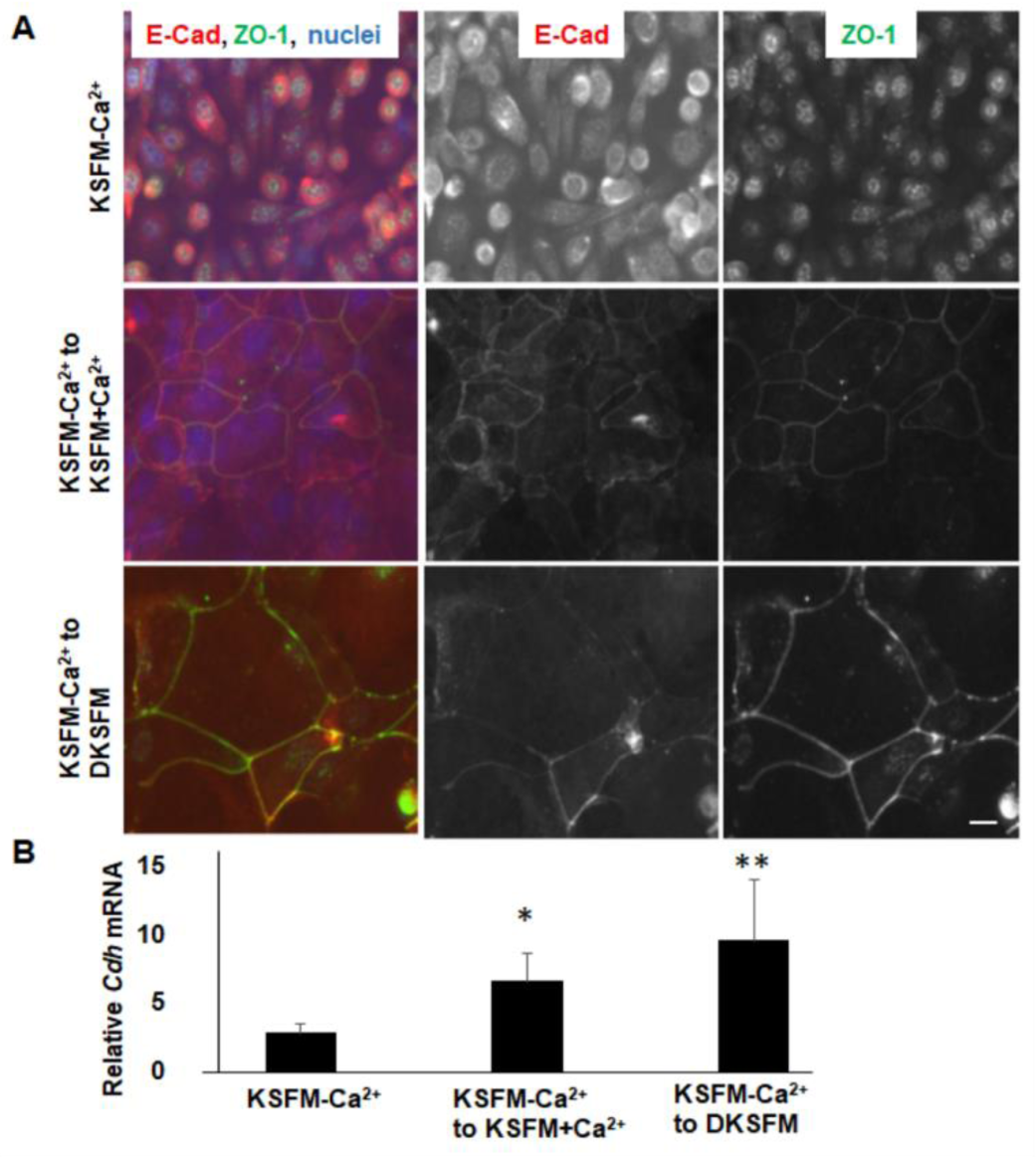
Restoration of cell-cell junctions in long-term cultured keratinocytes. Cells in KSFM-Ca^2+^ with or without switching to DKSFM or KSFM+Ca^2+^ for 48 h were examined by (A) immunofluorescence staining for E-Cadherin and ZO-1, and (B) RT-PCR for the expression of *Cdh* (N=3) +/-SD. Replacing the medium with DKSFM or KSFM+Ca^2+^ restored the formation of adherens junctions and tight junctions and significantly increased E-cadherin expression (*p<0.05, **p<0.01). Scale bar represents 20 µm.

### S1P activates the MAP3K1-JNK pathways

To assess gene functions in the keratinocytes, we made keratinocyte lines with KSFM-Ca^2+^ using cells isolated from wild type and *Map3k1*^*ΔKD*^ pups (13,53). The genotypic differences did not affect keratinocyte morphology or marker gene expression (Fig. S2).

We have previously shown, in mouse ESCs, that MAP3K1 is responsible for JNK activation by lysophosphatidic acid (LPA), which, like S1P, is a phospholipid signaling molecule that binds to and activates specific G-protein coupled receptors on the plasma membrane (24). To determine if MAP3K1 mediates the phospholipid signaling in keratinocytes, we treated the wild type and *Map3k1*^*ΔKD/ΔKD*^ cells with S1P and LPA and examined the phosphorylation of JNK, extracellular signal-regulated kinase (ERK) and p38 MAPKs. S1P and LPA induced a transient phosphorylation of JNK equally well in the wild type and the *Map3k1*^*ΔKD/ΔKD*^ cells grown in KSFM-Ca^2+^; however, they induced pJNK in the wild type but not the *Map3k1*^*ΔKD/ΔKD*^ cells that were switched to DKSFM medium for 48 h (Figs. 5A-C). Correspondingly, the phosphorylation of c-Jun, the JNK downstream substrate, was strongly induced by S1P and LPA in the wild type, but not in the *Map3k1*^*ΔKD/ΔKD*^ cells in DKSFM (Figs. 5B and 5D). The activation of ERK and p38, in contrast, was unaffected by *Map3k1* deficiency and the various growth medium conditions (Figs. 5B and 5C). Thus, MAP3K1 appears to be required for S1P-induced JNK activation only in the DKSFM medium-adapted keratinocytes.

**Figure 5.**
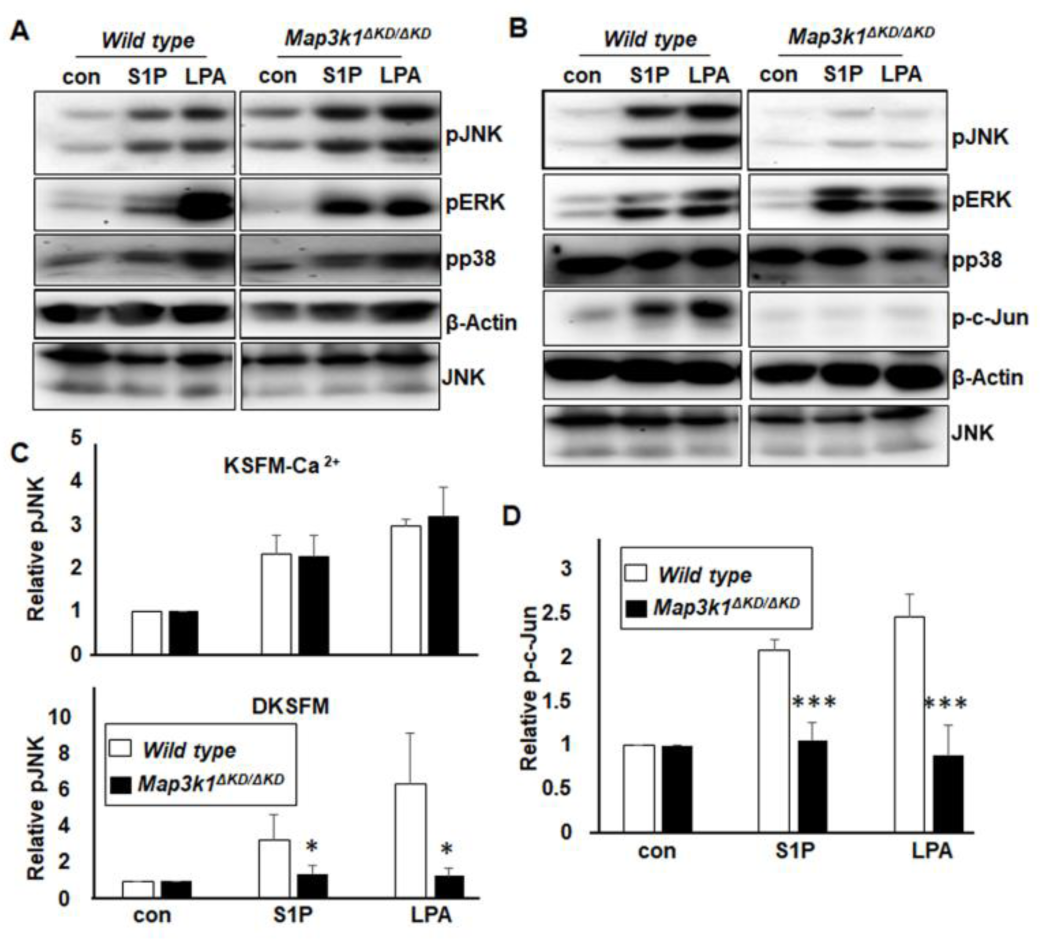
S1P activates the MAP3K1-JNK pathway in basal epithelial cells. The keratinocyte lines derived from wild type and *Map3k1*^*ΔKD/ΔKD*^ pups were either (A) maintained in KSFM-Ca^2+^ or (B) switched to DKSFM for 48 h. After starvation overnight in growth factor-free media, cells were treated with S1P (10 µM) and LPA (10 µM) for 30 min, and cell lysates analyzed by Western blotting using the antibodies indicated. The relative levels of (C) pJNK, and (D) p-c-Jun versus β-Actin were calculated based on signal intensity determined with the gel documentation system. In cells switched to DKSFM, the induction of pJNK and p-c-Jun was more abundant in wild type than in *Map3k1*^*ΔKD/ΔKD*^ cells. Results represent data from at least 5 independent experiments +/- SD *p<0.05, and ***p<0.001.

### The S1P-MAP3K1 pathway in epithelial cell movement and embryonic eyelid closure

The biological effects of S1P are mediated by the cell-surface G protein-coupled S1P receptors (31,33). Of the five receptor subtypes, S1PR1, 2 and 3 are expressed ubiquitously (54). Recent genetic data have shown that two of these receptors, S1PR2 and S1PR3, are collectively required for embryonic eyelid closure. Homozygous deletion of the gene encoding either receptor has no apparent effect on eyelid development, but the *S1pr2*^*-/-*^*S1pr3*^*-/-*^ pups are born with an EOB defect (36). While these findings suggest that S1PR2- and S1PR3-mediated signalling are required for eyelid closure, the downstream effector pathways remain elusive.

MAP3K1 and S1PR2/S1PR3 are expressed in the same group of epithelial cells in developing eyelids (13,36), leading us to interrogate their relationships *in vivo*. We crossed the *Map3k1*^+*/ΔKD*^ and the *S1pr2/S1pr3* mutant mice and examined the eyelid phenotypes in the newborn pups. While neither the *S1pr2*^+*/-*^*S1pr3*^*-/-*^ nor the *Map3k1*^+*/ΔKD*^ offspring had the open eye defects, approximately 50% of the *S1pr2*^+*/-*^*S1pr3*^*-/-*^*Map3k1*^+*/ΔKD*^ pups (N=6) displayed the EOB phenotype (Fig. 6A). We extended the genetic testing to evaluate the potential functional interplays between *S1pr* and *Jnk1*. In the *S1p2*^+*/-*^ *S1p3*^*-/-*^ genetic background, deletion of one *Jnk1* allele caused EOB defects in 5% pups (N =22), whereas ablation of two *Jnk1* alleles resulted in 63% of the pups (N=16) having the defects (Fig. 6A). The combined haplo-insufficiency of the mutants support the idea that S1PR2/3, MAP3K1 and JNK1 act in the same pathway for embryonic eyelid development (55).

**Figure 6.**
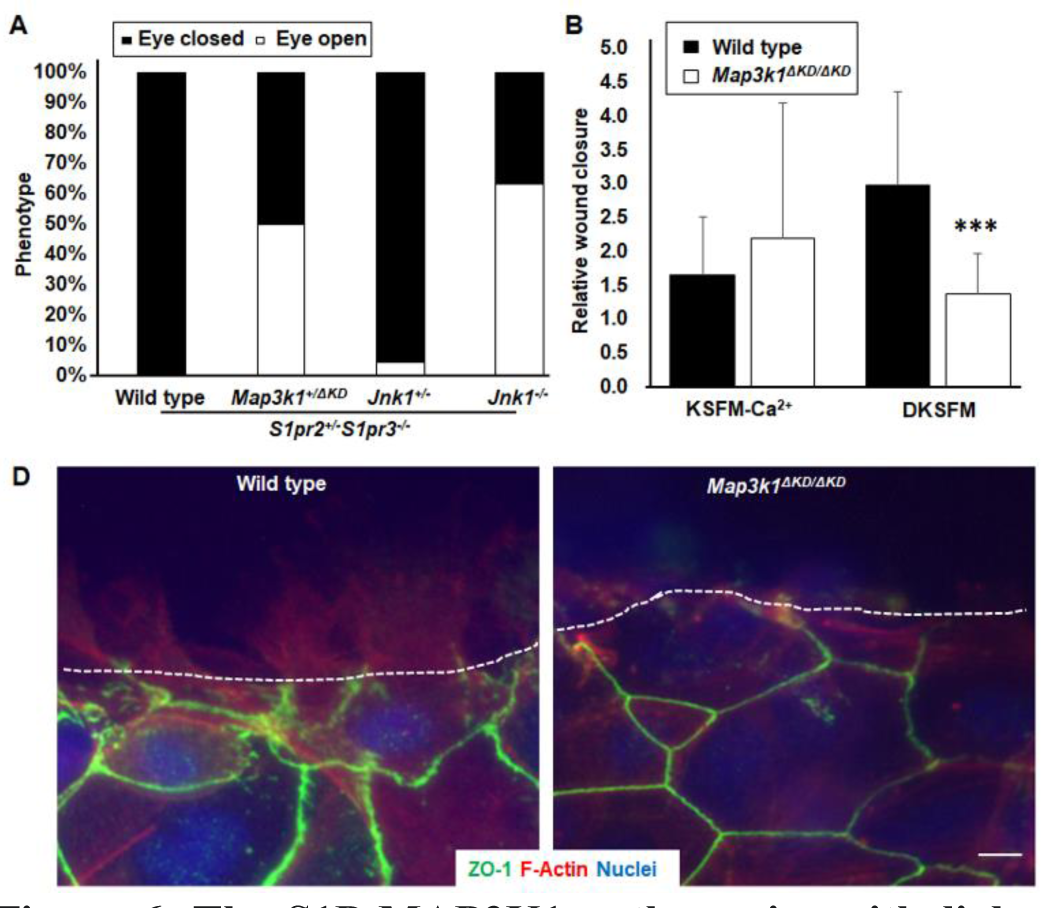
The S1P-MAP3K1 pathway in epithelial cell movement and eyelid closure. (A) Genetic crossing of the *S1p2/3*-mutant with the *Map3k1*- and the *Jnk1*-mutant mice. The eye open phenotype in the offspring of the indicated genotypes was recorded. A total of 19 *S1pr*^+*/-*^*S1pr3*^*-/-*^, 6 *S1pr2*^+*/-*^*S1pr3*^*-/-*^*Map3k1*^+*/ ΔKD*^, 22 *S1pr2*^+*/-*^*S1pr3*^*-/-*^ *Jnk1*^+*/-*^, and 16 *S1pr2*^+*/-*^*S1pr3*^*-/-*^ *Jnk1*^*-/-*^ pups were examined. (B) The keratinocyte lines derived from wild type and *Map3k1*^*ΔKD/ΔKD*^ pups were either maintained in KSFM-Ca^2+^ or switched to DKSFM for 48 h, and the confluent culture was subjected to *in vitro* wound-healing assays. The wounds were photographed at 0 and 24 h, when wound sizes were measured and wound-closure distances calculated. Results represent at least 6 independent experiments with 8 data sets in each experiment +/- SD. Compared to that of the wild type cells, the movement of the *Map3k1* ^*ΔKD*^ cells in the DKSFM was decreased *** *p*<0.001. (D) 6 h after wounding, keratinocytes grown in the DKSFM were fixed, permeabilized and subjected to immunofluorescence staining for ZO-1, phalloidin (F-actin) and Hoechst 33258 (nucleus). The wounding edge was photographed under the fluorescence microscope. Dotted lines mark the leading edge of the wound, scale bar represents 20 µm.

Embryonic eyelid closure is the result of morphogenetic movement of epithelial cells located at the developing eyelid margin (23). To examine if the S1P-MAP3K1 pathways were implicated in this movement, we performed the scratch wound healing assays with wild type and *Map3k1*^*ΔKD/ΔKD*^ keratinocytes. Addition of S1P to the wounded cells accelerated wound closure at similar rates, regardless of the *Map3k1* genotype, in cells grown in KSFM-Ca^2+^; however, it enhanced wound closure in wild type, but not *Map3k1*^*ΔKD*^ cells switched to the DKSFM (Fig. 6C). Correspondingly, the lamellipodia-like F-actin protrusion was detected at the leading edge of the wounded wild type cells, but was absent in that of *Map3k1*^*ΔKD/ΔKD*^ keratinocytes (Fig. 6D).

## Discussion

In this study, we identify the S1P/S1PR-MAP3K1 pathway as a key signaling mechanism in epithelial movement for eyelid morphogenesis. Genetic data showed that, whereas the *S1pr2*^*-/-*^*S1pr3*^*-/-*^ pups displayed the EOB phenotype, the *S1pr2*^+*/-*^*S1pr3*^*-/-*^ pups had normal eyelid development, suggesting a single *S1pr2* allele is sufficient for normal eyelid morphogenesis. The single *S1pr2* allele, however, became inadequate when the *Map3k1* was heterozygous, and up to 50% of the *S1pr2*^+*/-*^*S1pr3*^*-/-*^*Map3k1*^+*/ΔKD*^ pups exhibited the EOB defects. Similar to previous findings that *Jnk1*^*-/-*^ acts in synergy with *Map3k1*^+*/ΔKD*^ to cause the EOB phenotype (26), we found that *Jnk1* loss-of-function and *S1pr2*^+*/-*^*S1pr3*^*-/-*^ also exhibited haplo-insufficiency. Furthermore, *Jnk1* appears to contribute to eyelid closure signaling in a gene dose-dependent fashion, as loss of one *Jnk1* allele caused 5% EOB defects, and loss of two *Jnk1* alleles caused 63% EOB defects in the *S1pr2*^+*/-*^ *S1pr3*^*-/-*^ backgrounds. Together, the non-allelic non-complementation displayed by *S1pr2/3, Map3k1* and *Jnk1* suggests that these gene products functionally interact and contribute to the same biological pathway (55). Supporting the suggestion, *in vitro* studies in keratinocytes show that S1P is an extracellular cue that elicits a strong and rapid JNK activation mediated through MAP3K1. The S1P-induced MAP3K1-JNK pathway leads to accelerated epithelial movement, a cellular activity crucial for embryonic eyelid closure.

To date, long-term culture of mouse keratinocytes has remained a practical challenge. We report an experimental protocol for growing keratinocytes with extensive proliferative capacity. There are some procedures reported in the literature that involve multiple subcultures of murine keratinocytes (56-61). In comparison, our straightforward approach does not require a feeder layer nor conditioned medium or pre-selection and enrichment of a targeted subpopulation; it supports keratinocytes with exceedingly strong proliferation potential, allowing at least 60 passages. Keratinocytes grown under low-Ca^2+^ conditions, however, lacked adherens and tight junctions. When increased extracellular calcium, these cells restored cell-cell junctions and become responsive to S1P in activating the MAP3K1 pathway. The experimental approach described here enables the easy growth and subculture of keratinocytes derived from mice and promises an endless supply of genetically modified keratinocytes.

Of the three commercial keratinocyte media tested, only the KSFM-Ca^2+^ with 0.06 mM calcium supported long-term keratinocyte culture. *In vivo*, low Ca^2+^ concentrations are found in the inner basal layer of the epidermis, where epithelial cell proliferation and maintenance of homeostasis take place, whereas high Ca^2+^ concentrations are present in the outer suprabasal layers associated with increased keratinocyte terminal differentiation (62). The low Ca^2+^ conditions also seem to favor proliferation while prevent differentiation *in vitro*, because keratinocytes in KSFM-Ca^2+^ displayed increased proliferative markers, such Integrin α6 and p63, and decreased differentiation markers, such as K10. The proliferation capacity of the epidermis is attributed to progenitor, transient amplifying and stem cells, located in the interfollicular epidermis and the inner sheath of hair follicles (63-66). Our long-term cultured keratinocytes, however, did not have increased expression of selective stem cell markers, but had clonal expansion capacity. Given that the efficiency of generating the cell lines is relatively low, it is likely that the culture conditions described here selectively support a subset of epidermal progenitors, the molecular identities of which are yet to be fully understood.

As a phospholipid signaling molecule, S1P binds to and activates specific cell-surface GPCRs. The activated GPCRs in turn are coupled with heterotrimeric G-proteins to activate a range of downstream pathways, including AKT, RhoA, MAPK, and PLC (31). RhoA is a potential upstream activator of MAP3K1 because the active RhoA directly interacts with MAP3K1 (67) and RhoA loss-of-function delays eyelid closure in *Map3k1*^+*/ΔKD*^ pups by 2 days (27). Nevertheless, the *RhoA-null/Map3k1*^+*/ΔKD*^ eyelids eventually closed prenatally, and mutant pups were born with closed eyelids, suggesting RhoA makes a small contribution to MAP3K1 signaling. Moreover, we find that a chemical RhoA inhibitor only partially blocks S1P-induced JNK phosphorylation in keratinocytes (data not shown), suggesting other Rho family proteins, such as RhoB and RhoC, may play complementary roles in the transduction of S1P signals in MAP3K1 pathways.

Although LPA also activates the MAP3K1-JNK pathways in cultured keratinocytes, the LPA signaling appears to play little role in embryonic eyelid closure. LPA receptor null mice do not display an eyelid defect, and the presence of the LPA receptors does not rescue the EOB defects resulting from S1PR2/3 loss (68-71). It is reasonable to suggest that activation of the MAP3K1-JNK pathway by S1P is a developmental mechanism operative in a spatial-temporally specific fashion for eyelid morphogenetic closure (72). Whether this mechanism is implicated in other pathophysiological processes, such as tumor metastasis and inner ear development, where both the S1P receptors and the MAP3K1 play important roles, has become an intriguing open question (7,73-77).

A direct consequence of activating the S1P-MAP3K1-JNK pathway is the phosphorylation of c-Jun, a transcription factor of the activating protein-1 (AP-1) family. JNK-mediated c-Jun phosphorylation at serine 63 and 73 results in a significant increase of c-Jun transcription activity (78-80). c-Jun and its target genes, such as heparin-binding EGF (HB-EGF), are indeed implicated in epithelial migration and embryonic eyelid closure (81-84). However, c-Jun 63/73 (AA) mutant mice display normal eye development (85), suggesting c-Jun phosphorylation is not required for embryonic eyelid closure. Furthermore, *c-Jun* and *Map3k1* do not exhibit combined haploinsufficiency and their gene products are found in spatial-temporally separated cell populations in developing eyelids (86). On the other hand, JNK is known to carry out biological activities in a manner independent of c-Jun phosphorylation. For instance, JNK stimulates F-actin polymerization through the phosphorylation of paxillin, a focal adhesion-associated protein (87). This JNK activity leads to the formation of lamellipodia, major actin filament protrusions formed by migrating cells (88). In this context, we detected F-actin-enriched lamellipodia in S1P-treated wild type, not *Map3k1*^*ΔKD/ΔKD*^, migrating keratinocytes.

Eyelid closure defects lead to ocular adnexal structure abnormalities in post-natal life (89,90). The defective phenotypes resemble human congenital diseases, such as strabismus, ptosis and ectrodactyly-ectodermal dysplasia-cleft syndrome (91-94). The etiology and mechanisms for these diseases are still poorly understood. Data presented here suggest that abnormalities associated with defective eyelid closure could have polygenic etiology involving a combination of different genetic variants. Additionally, *in utero* exposure of the environmental toxicant dioxin induced the EOB phenotype in the *Map3k1*^+*/ΔKD*^ but not wild type pups, presenting a case where the eye defects were the result of gene-environment interactions and multifactorial etiology (95). It is thus reasonable to surmise that inactivation of the S1P-MAP3K1-JNK pathways is a molecular mechanism through which genetic and environmental factors induce defective eyelid closure and related developmental disorders.

### Experimental procedures

#### Experimental animals

Wild type and *Map3k1* ^*ΔKD*^ mice were backcrossed with BL6 mice for at least 10 generations, as described before (13,95). The *Map3k1*^*ΔKD/ΔKD*^ pups displayed the EOB phenotype and their genotypes were confirmed by established PCR methods. The *S1pr2* and *S1pr3* mutants were crossed and genotyped as previously described (54). Newborn pups were used for the isolation of primary keratinocytes using procedures approved by the University of Cincinnati Animal Care and Use Committee.

#### Chemicals, reagents and antibodies

The 3 keratinocyte culture media used were KSFM-Ca^2+^ (37010-022), KSFM+Ca^2+^ (17005-042) and DKSFM (10744019), from Gibco. TrypLE, 0.25% Trypsin without EDTA and the trypsin inhibitor were also from Gibco. CaCl_2_ solution was from PromoCell; Collagen IV was from BD Biosciences Discovery, and S1P and LPA were from Cayman Chemical. The antibodies for phospho-JNK and phospho-ERK were purchased from Cell Signaling; anti-phospho-p38 was from Promega; and the anti-β-Actin, calcium concentration kit, and Hoechst 33258 (for nuclei staining) were from Sigma. The anti-JNK antibody was obtained from Santa Cruz Biotechnology. Anti-ZO-1, anti-K14, Alexa fluor 594 phalloidin, anti-Rabbit Alexa Fluor 488 and anti-Mouse Alexa Fluor 594 were from Invitrogen, and anti-E-Cadherin was from BD Transduction Laboratories.

#### Mouse keratinocyte culture

Primary keratinocytes were isolated from newborn pups as described previously (38). Briefly, the epidermis was separated from dermis after overnight incubation in 0.25% trypsin, and placed in a tube containing 5 ml media with 1 mg/ml trypsin inhibitor. After shaking the tube 50 times, large epidermal pieces were removed and the cells were centrifuged at 2000 rpm for 5 min. The cells were resuspended in the medium of choice and plated on collagen IV-coated plates at >1×10^5^ cells/cm^2^. Cells were grown in a humidified CO_2_ incubator at 37 °C and refed every 2-3 days. Cells were passaged when reaching 90% confluence. For passaging, cells were rinsed with PBS, incubated with 0.05% EDTA in PBS for 10 min at 37 °C, trypsinized twice in TrypLE for 10 min at 37 °C, and collected in media containing 1 mg/ml trypsin inhibitor. Cells were centrifuged; the pellets were then resuspended in the medium of choice and plated on collagen IV-coated dishes. For routine passage, cells were plated at 1 - 2.5 x 10^4^/cm^2^.

For storage, cells dissociated from the culture dish were centrifuged as described above. The cell pellets were resuspended in 1 ml culture medium containing 10% DMSO and stored in liquid N_2_. The frozen cells were recovered after thawing in a 37 °C water bath, washed once with growth medium, centrifuged, and the pellets resuspended and plated in a collagen IV-coated dish.

In some experiments, when cells grown in KSFM-Ca^2+^ reached >80% confluence, the media were changed to either KSFM-Ca^2+^ plus 0.09 mM calcium, KSFM+Ca^2+^, or DKSFM, for 48 to 72 h.

#### Colony-forming efficiency (CFE) assays

Cells were plated at 1-200/cm^2^ and cultured for 14 days. After culture, the medium was removed, cells were fixed with 4 % PFA for 15 min, washed twice with PBS, and dried for 20 min. The cells were stained with 1% crystal violet at room temperature for 20 min; the plates were then rinsed with ample water and dried before photography and quantification.

#### Calcium concentration

The calcium concentrations in the growth media were determined using calcium concentration kit (Sigma), following the manufacturer’s protocol.

#### RNA isolation, reverse transcription and real-time quantitative polymerase chain reaction (RT-PCR)

Total RNA was isolated from cultured cells using PureLink RNA Mini Kit (Invitrogen). Reverse transcription was performed using SuperScript III reverse transcriptase (Invitrogen). RT-PCR was carried out with an Agilent Technologies Stratagene Mx3000P PCR machine using PowerUp SYBR Green Master Mix (Applied Biosystems) as the detection format. The reactions were cycled 40 times under the appropriate parameters for each pair of primers and fluorescence was measured at the end of each cycle to construct the amplification curve. All determinations were performed at least in triplicate. The primer sequences are included in Table S1.

#### Western blot analyses

The wild type and *Map3k1*^*ΔKD/ΔKD*^ keratinocytes maintained in KSFM-Ca^2+^ or switched to DKSFM for 2 days were starved overnight in corresponding media without growth factor (GF). Cells were treated with 10 uM S1P and 10 uM LPA in GF-free media for 30 min, and then lysed in RIPA buffer (150 mM NaCl, 1% Nonidet P-40, 0.5% sodium deoxycholate, 0.1% SDS, 50 mM Tris, pH 7.4). The cell lysates were subjected to SDS–PAGE, transferred onto nitrocellulose membranes and probed with antibodies as described previously (13).

#### Wound-healing assays

Wild type and *Map3k1*^*ΔKD/ΔKD*^ keratinocytes were grown in KSFM-Ca^2+^ in 24-well collagen IV-coated plates. When reaching 80% confluence, some cells were maintained in KSFM-Ca^2+^, and others had the medium changed to DKSFM for 24-48 hours. Scratch wounds were created on the monolayer with a micropipette tip and wound healing was carried out in GF-free KSFM-Ca^2+^ or DKSFM medium in the presence or absence of 10 µM S1P. At 24 hours after healing, the wound area was photographed, and the speed of wound closure was calculated based on wound area differences at 0 and 24 h. The wound closure rate was compared to that of untreated cells, defined as 1.

#### Immunofluorescence staining

Cells were seeded on collagen IV-coated 6 mm glass coverslips and grown in KSFM-Ca^2+^. When reaching 80% confluence, the medium was changed to DKSFM and/or KSFM + Ca^2+^ for 24-72 hours. In some experiments, scratch wounds were created and wounds were allowed to heal for 6 hours. The cells were fixed and permeabilized and immunostaining was performed as described previously (96). Images were obtained using a Zeiss Axio microscope.

#### Statistical Analyses

Means and standard deviations were calculated from at least three independent experiments, and analyzed using student’s t-test in which *p*<0.05 was considered statistically significant.

#### Data availability

These and all other data needed to evaluate the conclusions in the paper are present in the paper or the Supplementary Materials. All materials will be supplied upon request.

## Acknowledgements

The authors would like to thank Drs. Alvaro Pug, University of Cincinnati, for critical reading, Drs. Laura Wolszon and Gwendolyn Kaeser, Sanford Burnham Prebys Medical Discovery Institute, for editing, and Dr. Yuhang Zhang, University of Cincinnati, for consultations.

